# Brain Areas Affected by Intranasal Oxytocin Show Higher Oxytocin Receptor Expression

**DOI:** 10.1101/2021.01.12.426349

**Authors:** Philippe C Habets, Christabel Mclain, Onno C Meijer

## Abstract

Neuroimaging studies suggest that intranasal oxytocin (IN-OXT) may modulate emotional and social processes by altering neural activity patterns. The extent of brain penetration after IN-OXT is unclear, and it is currently under debate whether IN-OXT can directly bind central oxytocin receptors (OXTRs). We investigated oxytocin pathway gene expression in regions affected by IN-OXT on task-based fMRI. We found that OXTR is more highly expressed in affected than unaffected subcortical regions; this effect did not vary by task-type or sex. Cortical results revealed higher OXTR expression in regions affected by IN-OXT in emotional processing tasks and in male-only data. No significant differences were found in expression of the closely related vasopressin receptors. Our findings suggest that the mechanism by which IN-OXT may alter brain functionality involves direct activation of central OXTRs.

## Introduction

Oxytocin is a neuropeptide which has captured public and scientific interest in recent years due to its role in social behaviors and its potential as a novel treatment for neuropsychiatric disorders (1). This interest has given rise to many neuroimaging studies investigating the effects of intranasal oxytocin (IN-OXT) administration on brain activity (2). This body of research is contentious for two primary reasons: the neuroimaging results following IN-OXT administration have been inconsistent (2, 3), and the mechanism of action by which IN-OXT acts on the brain is unclear (4, 5). A recent study found that intranasal, but not intravenous, OXT administration in Macaques resulted in quantifiable exogenous OXT levels in multiple brain regions (6). This finding suggests that IN-OXT may bypass the blood-brain barrier to act directly on the brain, as has been posited before (7). However, penetrance was quite low and highly variable across animals in this study (6), casting additional doubt on the efficacy of IN-OXT delivery.

A direct mechanism of action would imply the ability of IN-OXT to bind to OXTRs in the brain. The distribution of receptors and binding patterns can be inferred from OXTR mRNA data. In addition to OXTR and OXT (the gene coding for the oxytocin prepropeptide), CD38 is a crucial oxytocin pathway gene which plays a role in oxytocin secretion and social behaviour (8). Expression of OXTR, OXT and CD38 is intercorrelated and highly variable throughout the subcortical human brain, with specific subcortical structures showing enrichment for these genes (including the hypothalamus) (9). Recent RNA sequencing datasets show low but variable expression of OXTR in multiple human cortical areas, while cortical expression of OXT and CD38 in the same RNA-seq datasets is hardly detectable with no measured variance (10, 11).

Here, we investigate expression of OXT, OXTR and CD38 in regions significantly affected by IN-OXT on task-based fMRI in humans, using microarray data from the Allen Human Brain Atlas (AHBA) (12). ‘Affected’ is defined as showing increased activity in IN-OXT over placebo conditions. We also investigate vasopressin receptor (AVPR) expression to control for possible IN-OXT-mediated fMRI effects via AVPR binding, as AVPRs have (weak) affinity for OXT and play an interrelated role in regulating social cognition and behavior (1, 13). Earlier work reported spatially correlated expression of OXTR and AVPRs (9). Accordingly, we assess correlations between all genes of interest (OXT, OXTR, CD38, AVPR1A, AVPR1B, AVPR2) to determine whether co-expression patterns differ in regions affected by IN-OXT on fMRI. We use statistic maps collated by Grace et al. which provide metadata from 39 studies, separated by task-type and sex (2). Using a stringent activation likelihood estimation (ALE) method, Grace et al. identified a cluster of convergence in experiments probing emotional, but not social, processes (2). Given the inconsistency in fMRI foci activated by IN-OXT and the poor penetrance via nasal delivery shown in animal models (6), we hypothesized that brain regions affected by IN-OXT on task-based fMRI would not show significantly higher expression of OXTR in comparison with unaffected brain regions.

## Results

### IN-OXT affected subcortical brain areas show higher expression of OXTR

Expression data for IN-OXT affected and unaffected subcortical and cortical samples in the emotion processing, social processing, combined and male-only masks are shown in Figure 1. For all four fMRI thresholded p-statistic maps, samples from IN-OXT affected subcortical brain regions showed significantly higher average expression of OXTR compared to unaffected subcortical brain regions. Oxytocin pathway genes OXT and CD38 were found to show a similar but weaker effect compared to OXTR, not reaching significance in all settings. No significant difference in subcortical expression was found for any of the AVP receptors.

**Figure 1.**
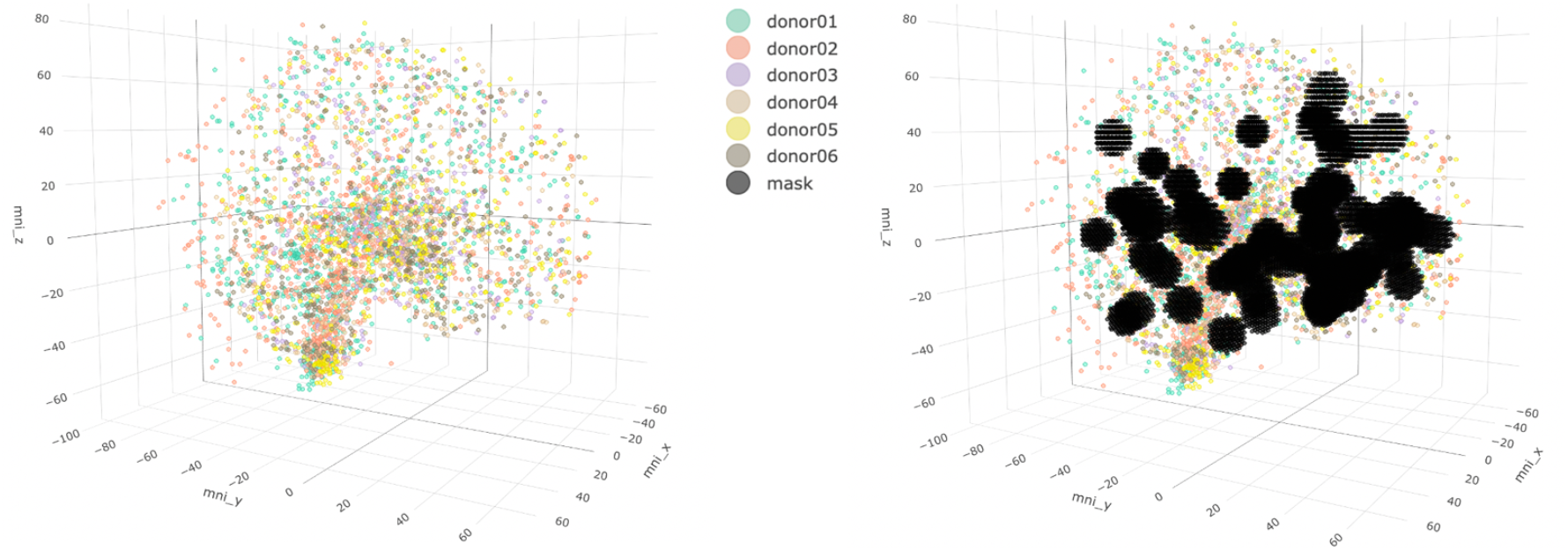
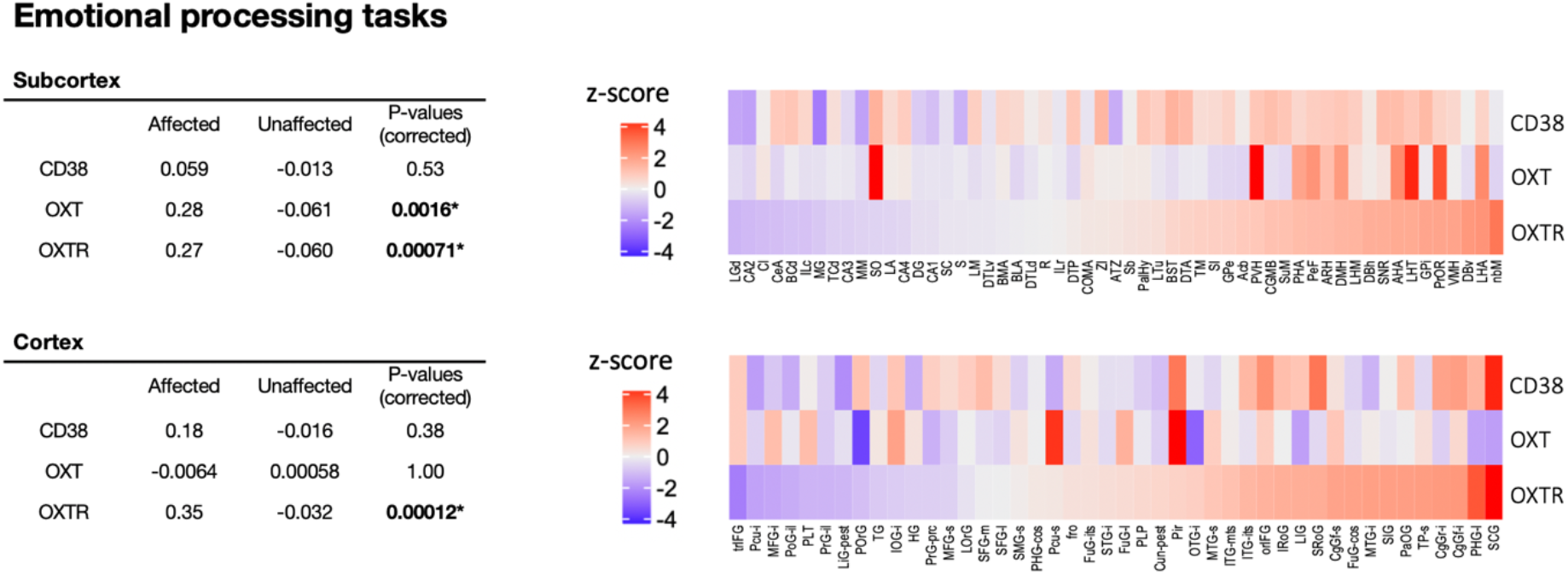
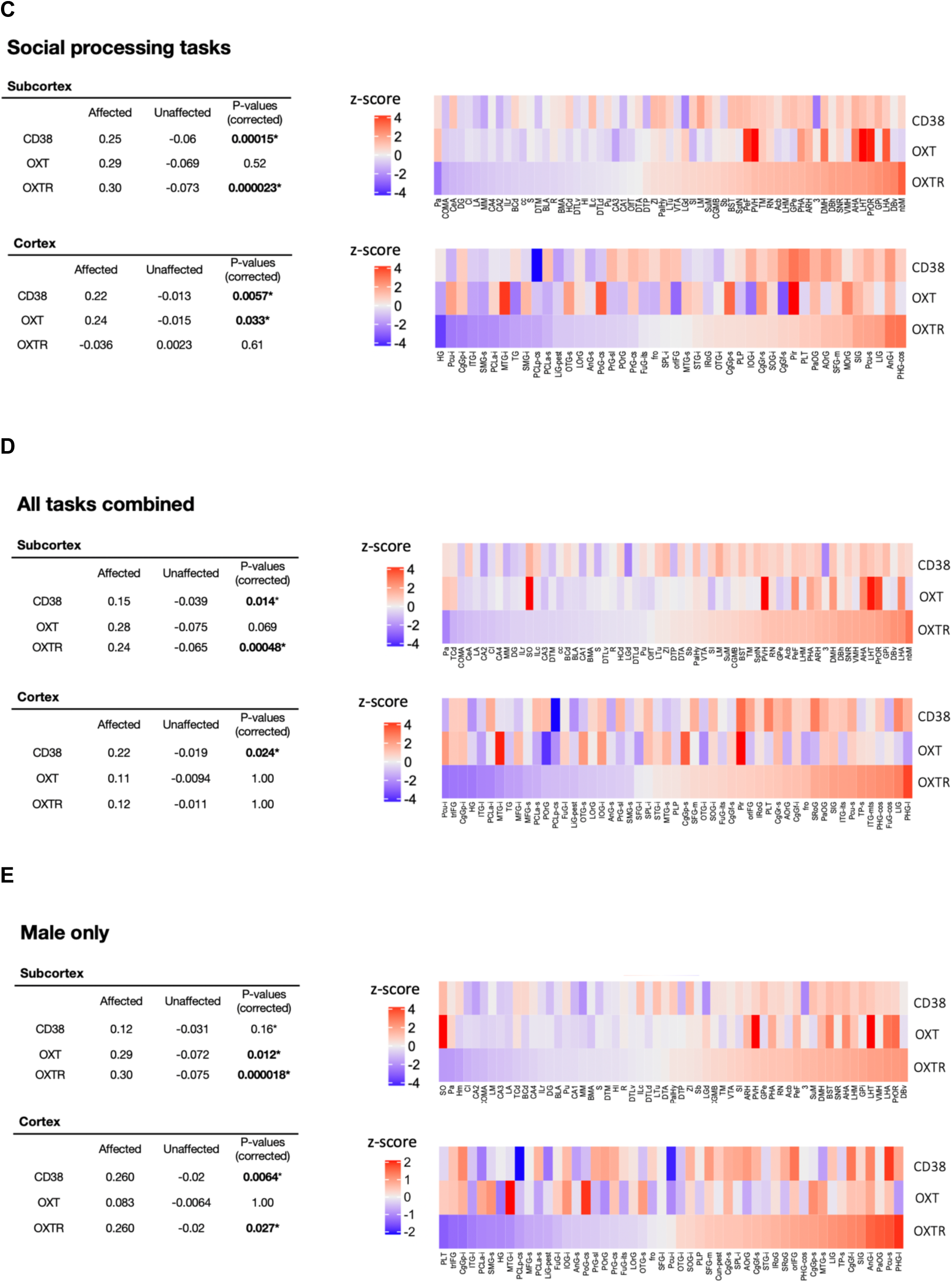
**A.** Brain samples from all six donors plotted in MNI-152 space (left). On the right, the thresholded p-statistic map for emotional processing is plotted in the same space. **B-E.** Differential gene expression analysis for CD38, OXT and OXTR in affected versus unaffected samples using four different p-statistic maps. Heatmaps show average expression of genes for brain structures that include affected samples. A legend of all brain structure abbreviations can be found in the supplemental data.

### IN-OXT affected cortical brain areas show differential expression for some OXT pathway genes

For cortical regions, areas affected by IN-OXT in emotional processing tasks showed significantly higher expression of OXTR. The same result was found for affected cortical regions in the male-only p-statistic map across all tasks. However, no significant difference in cortical OXTR expression was found in the other p-statistic maps (social processing and all combined OXT>PBO experiments). Except for the social processing p-statistic map, OXT showed no cortical differential expression in any of the included fMRI masks. With the exception of the emotional processing p-statistic map, a significantly higher expression of CD38 in affected cortical brain areas was found in all other masks. There was no significant difference in cortical expression of the AVP receptors in any of the fMRI statistic maps.

### IN-OXT affected subcortical brain areas show higher OXTR-AVPR1a co-expression

Pairwise gene correlations in affected samples (using the p-statistic map that combines all fMRI OXT>PBO experiments) are shown in Figure 2 for subcortex and cortex separately. Correlation matrices of subcortical affected versus unaffected samples differed significantly (χ2 = 53.55, p = 3.12e-6). Correlation matrices for the cortical samples showed no significant difference between affected versus unaffected samples (χ2 = 14.18, p = 0.51). Next, comparing OXTR co-expression in subcortical affected versus unaffected samples, a significant difference was found only for the co-expression of OXTR and AVPR1a (Table 2). OXTR and AVPR1a showed a significantly higher co-expression in the affected samples (r = 0.253 vs r = −0.00832, corrected p = 0.0026).

**Table 1.** Number of included samples for each group in the differential gene expression analysis.

|  |  | Emotion processing | Social processing | Male only | Overall |
| --- | --- | --- | --- | --- | --- |
| Subcortical samples | Affected | 177 | 193 | 176 | 208 |
|  | Unaffected | 809 | 793 | 705 | 778 |
| Cortical samples | Affected | 147 | 103 | 111 | 138 |
|  | Unaffected | 1615 | 1659 | 1427 | 1624 |

**Table 2.** Differences in correlation between subcortical affected and unaffected samples (all-task fMRI mask) for OXTR and other genes of interest. Δ-values (affected – unaffected correlation) are shown with Bonferroni corrected p-values computed by the Fisher’s z-test for correlation differences (two-tailed).

|  | **OXT** | **CD38** | **AVPR1A** | **AVPR1B** | **AVPR2** |
| --- | --- | --- | --- | --- | --- |
| **OXTR** | 0.17 (0.086) | 0.032 (<1) | 0.26 (0.0026)* | -0.010 (<1) | -0.12 (<1) |

**Figure 2.**
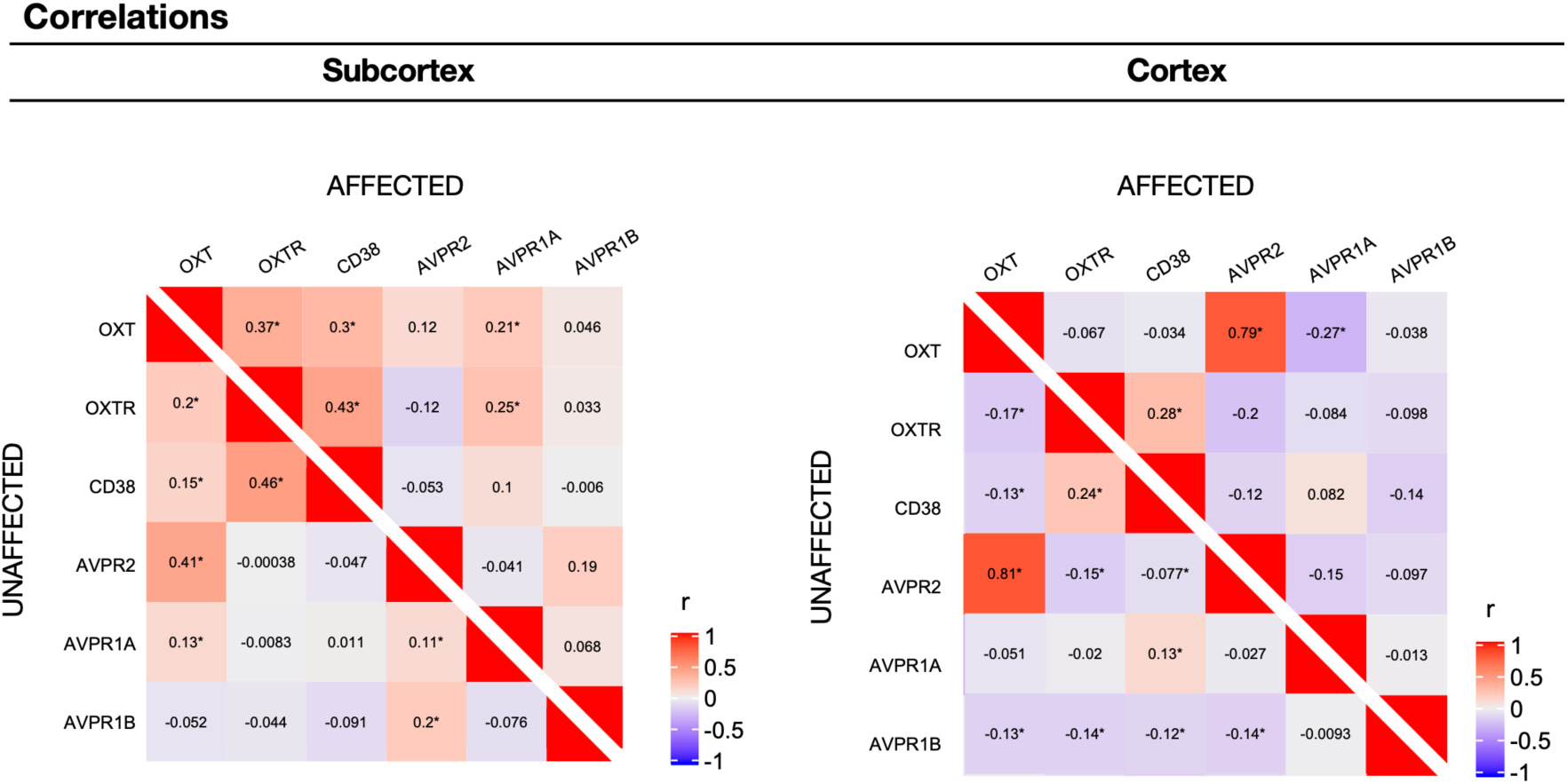
Pairwise correlations (Pearson’s r) of genes of interest in affected and unaffected samples, in subcortex and cortex respectively, using the all-task p-statistic map. Significant correlations (corrected p < 0.05) are marked with an asterisk (*).

## Discussion

Contrary to our hypothesis, we have identified robust differences in oxytocin pathway gene expression between IN-OXT-affected versus -unaffected subcortical brain regions. Remarkably, OXTR is more highly expressed in affected subcortical regions across all assessed fMRI files. As blood brain barrier penetration of IN-OXT is poor (5, 6), our findings support the hypothesis that IN-OXT acts directly on the brain via binding to its receptors in at least subcortical affected brain areas.

Cortical results were less consistent, with higher OXTR expression in affected areas on the emotion processing and male masks only. It is notable that the emotion processing file was the only mask for which the initial ALE meta-analysis by Grace et al. revealed a significant cluster of convergence (2). Tasks probing emotion were defined specifically as involving facial processing, while “social tasks” covered a much wider range (see Supplementary Table 3 of the report by Grace et al.) (2). This may have affected the specificity of the affected/unaffected regions on the social mask. Moreover, recent RNA-sequencing data indicate that cortical expression of OXTR is low, while OXT and CD38 is hardly detectable with no measured variance (11). Subcortical results thus provide stronger evidence from which to draw our conclusions. Although the thresholding of p-statistic maps is to some extent arbitrary, the risk of including false positive affected areas also comes with the risk of diluting any significant differential gene expression effect. It is therefore noteworthy that subcortical differential OXTR expression is found to be robust in all four fMRI masks.

OXTR and AVPR1a show significantly higher co-expression in affected versus unaffected subcortical brain areas. A higher correlation between these receptors does not necessarily indicate an interdependency between them (14). Rather, this co-expression may signify brain areas that have a function in multiple behavioral processes, some of which rely on OXTR and some on AVPRs. Thus, while IN-OXT mediated activity changes are specific to OXTR binding, affected brain areas likely play a role in other functions which may be sensitive to AVP.

The AHBA provides the most detailed dataset for examining spatial distribution of human brain transcriptomics to date but is limited to six donor brains. It is therefore important to note that independent sample validation of our genes of interest was performed in previous work using 10 overlapping brain regions from the Gentotype-Tissue Expression (GTEx) project (9, 15). OXTR and CD38, but not OXT, showed a significant correlation in expression between both datasets (9). Despite substantial differences in spatial coverage and donor number (for GTEx 88 to 173 donors are included depending on the brain region), brain co-expression patterns of oxytocin pathway genes were found to be similar for the comparable 10 brain regions (details described in Quintana et al.) (9). This lends validity to our use of the AHBA for analysis.

By providing evidence for a hypothesized mechanism of action, our findings serve to attenuate skepticism towards the ability of IN-OXT to affect brain functionality.

## Materials and Methods

SI Materials and Methods including detailed description of our differential expression and correlation analyses are given in the supplementary methods section. All code and used files are made publicly available on GitHub: https://github.com/pchabets/fMRI-Transcriptomics-Oxytocin.

## References

1. A. Meyer-Lindenberg, G. Domes, P. Kirsch, M. Heinrichs, Oxytocin and vasopressin in the human brain: social neuropeptides for translational medicine. Nat. Rev. Neurosci. 12, 524–538 (2011).

2. S. A. Grace, S. L. Rossell, M. Heinrichs, C. Kordsachia, I. Labuschagne, Oxytocin and brain activity in humans: A systematic review and coordinate-based meta-analysis of functional MRI studies. Psychoneuroendocrinology 96, 6–24 (2018).

3. D. S. Quintana, Revisiting non-significant effects of intranasal oxytocin using equivalence testing. Psychoneuroendocrinology 87, 127–130 (2018).

4. G. Leng, M. Ludwig, Intranasal Oxytocin: Myths and Delusions. Biol. Psychiatry 79, 243– 250 (2016).

5. D. S. Quintana, K. T. Smerud, O. A. Andreassen, P. G. Djupesland, Evidence for intranasal oxytocin delivery to the brain: recent advances and future perspectives. Ther. Deliv. 9, 515–525 (2018).

6. M. R. Lee, et al., Labeled oxytocin administered via the intranasal route reaches the brain in rhesus macaques. Nat. Commun. 11, 2783 (2020).

7. F. Erdő, L. A. Bors, D. Farkas, Á. Bajza, S. Gizurarson, Evaluation of intranasal delivery route of drug administration for brain targeting. Brain Res. Bull. 143, 155–170 (2018).

8. D. Jin, et al., CD38 is critical for social behaviour by regulating oxytocin secretion. Nature 446, 41–45 (2007).

9. D. S. Quintana, et al., Oxytocin pathway gene networks in the human brain. Nat. Commun. 10, 668 (2019).

10. R. D. Hodge, et al., Conserved cell types with divergent features in human versus mouse cortex. Nature 573, 61–68 (2019).

11. Allen Institute for Brain Science. Allen Cell Types Database (2019). Available at https://celltypes.brain-map.org/rnaseq/human_ctx_smart-seq.

12. M. J. Hawrylycz, et al., An anatomically comprehensive atlas of the adult human brain transcriptome. Nature 489, 391–399 (2012).

13. M. Heinrichs, G. Domes, Neuropeptides and social behaviour: effects of oxytocin and vasopressin in humans. Prog. Brain Res. 170, 337–350 (2008).

14. A. Galbusera, et al., Intranasal Oxytocin and Vasopressin Modulate Divergent Brainwide Functional Substrates. Neuropsychopharmacology 42, 1420–1434 (2017).

15. Human genomics. The Genotype-Tissue Expression (GTEx) pilot analysis: multitissue gene regulation in humans. Science 348, 648–660 (2015).

