## Extended Methods and Data for "Brain Areas Affected by Intranasal Oxytocin Show Higher Oxytocin Receptor Expression"

**Supplementary Methods.**

***Transcriptomic atlas***

The Allen Human Brain Atlas (AHBA) is a publicly available transcriptional atlas based on microarray measures in 3702 samples across brainstem, cerebellum, subcortical and cortical brain structures across six postmortem human brains (five male and one female). For limited samples of two donor brains, expression values were also measured by RNA sequencing. All expression data and metadata were downloaded from the AHBA (http://human.brain-map.org) (1, 2).

***fMRI data***

Data from 39 selected IN-OXT fMRI studies were included in the study by Grace et al (3). For our analysis, we looked at four different p-statistic maps. Each statistic map was created separately for differential activation in the intranasal-oxytocin versus placebo conditions (OXT>PBO) in:

1) tasks related to emotional processing (14 experiments, 72 foci, 506 participants)
2) tasks related to social processing (16 experiments, 153 foci, 873 participants)
3) all experiments that looked at the OXT>PBO contrast (31 experiments, 243 foci, 1420 participants)
4) all experiments that looked at the OXT>PBO contrast but including data from male participants only (22 experiments, 142 foci, 855 participants)

Details about study selection, ALE analysis and statistic map generation is found in the original paper by Grace et al (3).

All four p-statistic maps were downloaded from Neurovault at https://neurovault.org/collections/3713/ (map ID: 63321, 63369, 63357, 63349). Each statistic map was then thresholded for a p-value of 0.05. The resulting thresholded foci were considered as "IN-OXT-affected brain areas", and assessed for differential gene expression in comparison to other brain areas that, after thresholding, were considered "IN-OXT-unaffected".

***Data analysis***

With the exception of probe reannotation, R was used for all data analysis, with all used packages installed under R version 4.0.

First, we re-annotated all microarray probes for the genes of interest to the latest human genome version and reference sequence (May 20th, 2020), using the Re-Annotator package (4). Re-Annotator is freely available for download at https://sourceforge.net/projects/reannotator/.

Next, given that more than one probe was annotated to each gene of interest (OXT, OXTR, CD38, AVPR1A, AVPR1B, AVPR2), we selected a probe for each gene on the basis of highest expression (intensity analysis). We validated probe selections by confirming that each probe also showed the highest correlation (Pearson's r) with the RNA sequencing measures for the same gene, from the same brain sample (data only available for the first two donor brains) (5).

AHBA samples and fMRI masks were plotted in MNI-152 space**.** Using trilinear interpolation, we calculated whether each AHBA sample could be assigned to an affected brain area (meaning falling inside the fMRI mask) or not. Brainstem and cerebellum samples were excluded from analysis. Because the rate of 'affected' versus 'unaffected' samples differed between cortical and subcortical samples for all statistic maps, we separated cortical and subcortical samples in our analyses to prevent bias by the inherently different gene expression profiles of the cortex and subcortex (2). This resulted in the selection of samples that were categorized in one of four groups:

1. subcortical IN-OXT unaffected samples;
2. subcortical IN-OXT affected samples;
3. cortical IN-OXT unaffected samples;
4. cortical IN-OXT affected samples.

We corrected downloaded expression values from the AHBA for between-donor differences that might drive differential gene expression findings by using the *removeBatchEffect()* function from the limma package for R (6), treating each donor as a separate batch. It is relevant to note that gene expression patterning across brain structures was assessed for reproducibility in all six AHBA donor brains in previous works (2, 7). This was done using the differential stability (DS) metric: a measure for the consistency of a gene's differential expression pattern across brain structures (8). In one study that included 17,348 protein coding genes in the DS analysis, all oxytocin pathway genes showed a DS of 0.65 or higher, with OXTR and OXT in the top 10% of genes with highest DS scores (see supplementary table 2 of Hawrylycz et al.) (2). Another study by Quintana et al. found OXTR and CD38 to both have top decile DS scores in a list of 20,737 protein coding genes (7). These results suggests that, at least in the six donor brains, oxytocin pathway genes show gene expression patterning that is reproducible across donor brains, regardless of individual differences like sex or age.

Next, we z-normalized expression values across brains for use in heatmap visualization and further assessment. Data from all six donor brains were used for assessing differential gene expression in affected versus unaffected brain areas in the first three p-statistic maps. For the p-statistic map that includes male participants only, expression data from only the five male donor brains were used.

We assessed differential gene expression between affected and unaffected samples separately for subcortical and cortical samples using a Wilcoxon-Rank Sum test. We controlled for family-wise error rates by using Bonferroni correction of p-values. For all our analyses we used Bonferroni-corrected $p < 0.05$ as an indication of significance. All steps after probe selection were repeated for the four different fMRI masks.

We assessed correlations between all genes of interest in IN-OXT-affected and -unaffected brain areas, using the p-statistic map of all experiments that looked at the OXT>PBO contrast. We calculated Pearson' for both groups of samples (subcortical affected versus unaffected, cortical affected versus unaffected respectively), incorporating expression data from all six donor brains. This resulted in four correlation matrices. Pairs of correlation matrices corresponding to affected and unaffected samples in the subcortical and cortical subgroups were respectively tested for equality, using the *cortest.normal()* function of the 'psych' package for R (this function uses the Steiger test for comparing correlation matrices) (9).

After differences in overall gene correlation matrices had been assessed (only significant for subcortical groups, see *Results*), we tested for OXTR co-expression differences (i.e., differences in Pearson's r) in the subcortical affected versus unaffected samples. We first excluded insignificant gene correlation coefficients (Bonferroni-corrected $p >= 0.05$) in both affected versus unaffected samples. We assessed significance in differences between the remaining correlation coefficients for OXTR and the other genes of interest using Fisher's Z-transformation test (two-tailed, Bonferroni-corrected $p < 0.05$).

**Extended Methods References**


1. M. J. Hawrylycz, *et al.*, An anatomically comprehensive atlas of the adult human brain transcriptome. *Nature* **489**, 391–399 (2012).
2. M. Hawrylycz, *et al.*, Canonical genetic signatures of the adult human brain. *Nat. Neurosci.* **18**, 1832–1844 (2015).
3. S. A. Grace, S. L. Rossell, M. Heinrichs, C. Kordsachia, I. Labuschagne, Oxytocin and brain activity in humans: A systematic review and coordinate-based meta-analysis of functional MRI studies. *Psychoneuroendocrinology* **96**, 6–24 (2018).
4. J. Arloth, D. M. Bader, S. Röh, A. Altmann, Re-Annotator: Annotation Pipeline for Microarray Probe Sequences. *PLoS One* **10**, e0139516 (2015).
5. A. Arnatkevic Iute, B. D. Fulcher, A. Fornito, A practical guide to linking brain-wide gene expression and neuroimaging data. *Neuroimage* **189**, 353–367 (2019).
6. M. E. Ritchie, *et al.*, limma powers differential expression analyses for RNA-sequencing and microarray studies. *Nucleic Acids Res.* **43**, e47–e47 (2015).
7. D. S. Quintana, *et al.*, Oxytocin pathway gene networks in the human brain. *Nat. Commun.* **10**, 668 (2019).
8. G. T. W. Shaw, E. S. C. Shih, C.-H. Chen, M.-J. Hwang, Preservation of Ranking Order in the Expression of Human Housekeeping Genes. *PLoS One* **6**, e29314 (2011).
9. W. Revelle, psych: Procedures for Psychological, Psychometric, and Personality Research (2020).

**Supplemental Data**

Structure acronyms (as they appear in Figure 1) and their corresponding names, extracted from the AHBA sample information for all six donor brains.

| Stucture Acronym | Structure Name |
|---|---|
| PCLa-i | paracentral lobule, anterior part, right, inferior bank of gyrus |
| Cl | claustrum, right |
| LGd | dorsal lateral geniculate nucleus, left |
| CA4 | CA4 field, right |
| DG | dentate gyrus, right |
| Dt | dentate nucleus, left |
| Fas | fastigial nucleus, left |
| Emb | emboliform nucleus, right |
| S | subiculum, left |
| CA1 | CA1 field, left |
| TCd | tail of caudate nucleus, left |
| DTA | anterior group of nuclei, right |
| DTLv | lateral group of nuclei, right, ventral division |
| ILr | rostral group of intralaminar nuclei, right |
| DTM | medial group of nuclei, left |
| ILc | caudal group of intralaminar nuclei, left |
| R | reticular nucleus of thalamus, right |
| Pa | paraventricular nuclei, right of thalamus, right |
| SI | substantia innominata, right |
| Sb | subthalamic nucleus, right |
| SNC | substantia nigra, pars compacta, left |
| ZI | zona incerta, left |
| CGMB | central gray substance of midbrain, right |
| RN | red nucleus, left |
| SNR | substantia nigra, pars reticulata, left |
| MTG-i | middle temporal gyrus, left, inferior bank of gyrus |
| ATZ | amygdalohippocampal transition zone, left |
| BLA | basolateral nucleus, left |
| BMA | basomedial nucleus, left |
| LA | lateral nucleus, left |
| CeA | central nucleus, left |
| SIG | short insular gyri, left |
| PoG-cs | postcentral gyrus, right, bank of the central sulcus |
| OTG-i | occipito-temporal gyrus, left, inferior bank of gyrus |
| FuG-its | fusiform gyrus, left, bank of the its |
| FuG-l | fusiform gyrus, left, lateral bank of gyrus |
| FuG-cos | fusiform gyrus, left, bank of cos |
| HG | Heschl's gyrus, left |
| Pu | putamen, left |
| LIG | long insular gyri, right |
| STG-i | superior temporal gyrus, left, inferior bank of gyrus |
| MTG-s | middle temporal gyrus, left, superior bank of gyrus |
| ITG-l | inferior temporal gyrus, left, lateral bank of gyrus |
| ITG-mts | inferior temporal gyrus, left, bank of mts |
| STG-l | superior temporal gyrus, right, lateral bank of gyrus |
| ITG-its | inferior temporal gyrus, right, bank of the its |
| PoG-il | postcentral gyrus, right, inferior lateral aspect of gyrus |
| PLT | planum temporale, right |
| PrG-prc | precentral gyrus, left, bank of the precentral sulcus |
| PrG-sl | precentral gyrus, left, superior lateral aspect of gyrus |

| | |
|---|---|
| PrG-il | precentral gyrus, left, inferior lateral aspect of gyrus |
| MFG-i | middle frontal gyrus, left, inferior bank of gyrus |
| SFG-m | superior frontal gyrus, left, medial bank of gyrus |
| SFG-l | superior frontal gyrus, left, lateral bank of gyrus |
| MFG-s | middle frontal gyrus, left, superior bank of gyrus |
| PrG-cs | precentral gyrus, right, bank of the central sulcus |
| GPi | globus pallidus, internal segment, right |
| PHG-l | parahippocampal gyrus, left, lateral bank of gyrus |
| PHG-cos | parahippocampal gyrus, left, bank of the cos |
| CgGp-s | cingulate gyrus, parietal part, left, superior bank of gyrus |
| CgGp-i | cingulate gyrus, parietal part, left, inferior bank of gyrus |
| cc | corpus callosum |
| cgb | cingulum bundle, right |
| PLP | planum polare, left |
| SMG-s | supramarginal gyrus, left, superior bank of gyrus |
| BCd | body of caudate nucleus, right |
| SMG-i | supramarginal gyrus, left, inferior bank of gyrus |
| CgGf-s | cingulate gyrus, frontal part, left, superior bank of gyrus |
| CgGf-i | cingulate gyrus, frontal part, left, inferior bank of gyrus |
| PoG-sl | postcentral gyrus, right, superior lateral aspect of gyrus |
| GPe | globus pallidus, external segment, right |
| TG | transverse gyri, right |
| PoG-pcs | postcentral gyrus, left, bank of the posterior central sulcus |
| PCLa-s | paracentral lobule, anterior part, right, superior bank of gyrus |
| AnG-i | angular gyrus, left, inferior bank of gyrus |
| SPL-i | superior parietal lobule, left, inferior bank of gyrus |
| HCd | head of caudate nucleus, right |
| AOrG | anterior orbital gyrus, right |
| LOrG | lateral orbital gyrus, right |
| orIFG | inferior frontal gyrus, orbital part, left |
| GRe | gyrus rectus, right |
| IRoG | inferior rostral gyrus, left |
| SRoG | superior rostral gyrus, right |
| OTG-s | occipito-temporal gyrus, left, superior bank of gyrus |
| LiG-pest | lingual gyrus, right, peristriate |
| LiG-str | lingual gyrus, right, striate |
| Cun-pest | cuneus, right, peristriate |
| SOG-s | superior occipital gyrus, left, superior bank of gyrus |
| AnG-s | angular gyrus, right, superior bank of gyrus |
| Pcu-i | precuneus, right, inferior lateral bank of gyrus |
| SPL-s | superior parietal lobule, right, superior bank of gyrus |
| Pcu-s | precuneus, right, superior lateral bank of gyrus |
| MOrG | medial orbital gyrus, left |
| Cun-str | cuneus, left, striate |
| FP-s | frontal pole, left, superior aspect |
| CA2 | CA2 field, right |
| CA3 | CA3 field, right |
| LHM | lateral hypothalamic area, mammillary region, left |
| PHA | posterior hypothalamic area, left |
| VTA | ventral tegmental area, left |
| 3 | oculomotor nuclear complex, left |
| MPB | medial parabrachial nucleus, left |
| Pr5 | principal sensory nucleus of trigeminal nerve, left |
| 7 | facial motor nucleus, left |
| 12 | hypoglossal nucleus, right |
| COMA | cortico-medial group, left |
| RaM | raphe nuclei of medulla |
| Mo5 | motor nucleus of trigeminal nerve, right |
| IO | inferior olivary complex, right |
| LPB | lateral parabrachial nucleus, left |

| | |
|---|---|
| LC | locus ceruleus, right |
| Pn | pontine nuclei, right |
| SubC | nucleus subceruleus, left |
| PRF | pontine reticular formation, left |
| Arc | arcuate nucleus of medulla, right |
| MBRF | midbrain reticular formation, left |
| SubCn | subcuneiform nucleus, right |
| SOC | superior olivary complex, left |
| 4 | trochlear nucleus, right |
| MBRa | midbrain raphe nuclei |
| fro | frontal operculum, right |
| opIFG | inferior frontal gyrus, opercular part, right |
| Ve-VIIAt | VIIAt |
| PV-Crus I | Crus I, left, paravermis |
| PV-VIIIA | VIIIA, right, paravermis |
| MG | medial geniculate complex, left |
| Glo | globose nucleus, left |
| DTLd | lateral group of nuclei, left, dorsal division |
| DTP | posterior group of nuclei, left |
| SC | superior colliculus, left |
| SptN | septal nuclei, left |
| PrOR | preoptic region, left |
| SO | supraoptic nucleus, left |
| PTec | pretectal region |
| Hm | medial habenular nucleus, right |
| Hl | lateral habenular nucleus, right |
| PaOG | parolfactory gyri, left |
| SCG | subcallosal cingulate gyrus, right |
| GiRt | gigantocellular group, left |
| LMRt | lateral medullary reticular group, right |
| Sp5 | spinal trigeminal nucleus, left |
| 8Ve | vestibular nuclei, left |
| 10 | dorsal motor nucleus of the vagus, left |
| 6 | abducens nucleus, left |
| RPn | pontine raphe nucleus |
| 8Co | cochlear nuclei, right |
| trIFG | inferior frontal gyrus, triangular part, left |
| POrG | posterior orbital gyrus, right |
| TP-m | temporal pole, right, medial aspect |
| TP-s | temporal pole, right, superior aspect |
| TP-i | temporal pole, right, inferior aspect |
| CGS | central glial substance |
| Cu | cuneate nucleus, left |
| CMRt | central medullary reticular group, left |
| Gr | gracile nucleus, right |
| PCLp-cs | paracentral lobule, posterior part, right, bank of cingulate sulcus |
| PCLp-l | paracentral lobule, posterior part, right, lateral bank of gyrus |
| PV-VI | VI, right, paravermis |
| Acb | nucleus accumbens, left |
| He-VI | VI, left, lateral hemisphere |
| He-Crus I | Crus I, left, lateral hemisphere |
| He-Crus II | Crus II, left, lateral hemisphere |
| He-VIIB | VIIB, left, lateral hemisphere |
| He-VIIIA | VIIIA, left, lateral hemisphere |
| Ve-I-II | I-II |
| Ve-III | III |
| Ve-IX | IX |
| Ve-IV | IV |
| Ve-V | V |

| | |
|---|---|
| Ve-VI | VI |
| Ve-VIIAf | VIIAf |
| Ve-VIIB | VIIB |
| Ve-VIIIA | VIIIA |
| Ve-VIIIB | VIIIB |
| PV-III | III, left, paravermis |
| PV-IV | IV, left, paravermis |
| PV-V | V, left, paravermis |
| PV-VIIIB | VIIIB, left, paravermis |
| PV-IX | IX, left, paravermis |
| PV-Crus II | Crus II, left, paravermis |
| PV-VIIB | VIIB, left, paravermis |
| FPi | frontal pole, left, inferior aspect |
| FPm | frontal pole, left, medial aspect |
| Ve-X | X |
| PV-X | X, left, paravermis |
| CgGr-s | cingulate gyrus, retrosplenial part, right, superior bank of gyrus |
| CgGr-i | cingulate gyrus, retrosplenial part, right, inferior bank of gyrus |
| PTG | paraterminal gyrus, right |
| IOG-s | inferior occipital gyrus, right, superior bank of gyrus |
| SOG-i | superior occipital gyrus, right, inferior bank of gyrus |
| IOG-i | inferior occipital gyrus, right, inferior bank of gyrus |
| CGPo | central gray of the pons, left |
| EW | Edinger-Westphal nucleus, right |
| Dk | nucleus of Darkschewitsch, left |
| ICjl | interstitial nucleus of Cajal, right |
| IC | inferior colliculus, left |
| CnF | cuneiform nucleus, right |
| ARH | arcuate nucleus of the hypothalamus, left |
| PVH | paraventricular nucleus of the hypothalamus, left |
| LHA | lateral hypothalamic area, anterior region, left |
| AHA | anterior hypothalamic area, left |
| VMH | ventromedial hypothalamic nucleus, left |
| MB | mammillary body, left |
| PCLa | paracentral lobule, anterior part, left |
| LTu | lateral tuberal nucleus, left |
| PeF | perifornical nucleus, left |
| nbM | basal nucleus of meynert, left |
| DBh | nucleus of the diagonal band, left, vertical division |
| DBv | nucleus of the diagonal band, left, horizontal division |
| CPLV | choroid plexus of the lateral ventricle |
| SuM | supramammillary nucleus, left |
| LM | lateral mammillary nucleus, left |
| MM | medial mammillary nucleus, left |
| LHT | lateral hypothalamic area, tuberal region, left |
| TM | tuberomammillary nucleus, left |
| DMH | dorsomedial hypothalamic nucleus, left |
| BST | bed nucleus of stria terminalis, left |
| OlfT | olfactory tubercle, left |
| He-IX | IX, left, lateral hemisphere |
| He-V | V, left, lateral hemisphere |
| He-IV | IV, left, lateral hemisphere |
| He-III | III, left, lateral hemisphere |
| He-VIIIB | VIIIB, left, lateral hemisphere |
| PIN | pineal gland |
| Pir | piriform cortex, left |
| PalHy | pallidohypothalamic nucleus, left |
| He-X | X, left, lateral hemisphere |
